# Rigor and Reproducibility of Digital Spatial Profiling on Clinically Sourced Human Tissues

**DOI:** 10.1101/2024.10.16.618750

**Authors:** Kelly D. Smith, James W. MacDonald, Xianwu Li, Emily Beirne, Galen Stewart, Theo K. Bammler, Shreeram Akilesh

**Affiliations:** Department of Laboratory Medicine and Pathology, University of Washington, Seattle, WA 98195; Department of Environmental and Occupational Health Sciences, University of Washington, Seattle, WA 98105

**Author notes:** Corresponding authors Kelly D. Smith, Shreeram Akilesh, 1959 NE Pacific St, Box 356100, Seattle, WA 98195. Equal contribution (Kelly D. Smith, Shreeram Akilesh).

**Keywords:** Spatial transcriptomics, rigor and reproducibility, GeoMx DSP, CosMx SMI, human biopsy tissue, digital spatial profiling, single molecular imaging

## Abstract

Spatial transcriptomic profiling enables precise quantification of gene expression with simultaneous localization of expression profiles onto tissue structures. This new technology promises to improve our understanding of the disease mechanisms. Therefore, there is intense interest in applying these methods in clinical trials or as laboratory developed tests to aid in diagnosis of disease. Before these technologies can be more broadly deployed in clinical research and diagnostics, it is necessary to thoroughly understand their performance in real world conditions. In this study, we vet technical reproducibility, data normalization methods and assay sensitivity focusing predominantly on one widely used spatial transcriptomic methodology, digital spatial profiling. Using clinically sourced human tissue specimens, we find that digital spatial profiling exhibits high rigor and reproducibility. Our approach lays the foundation for incorporation of digital spatial profiling methods into clinical workflows.

## INTRODUCTION

Tissues have complex architecture with multiple cell types and cell states. Traditional approaches to analyze bulk gene expression and single cell gene expression destroy the complex architecture of tissues and the relationships between cells and extracellular components. Spatial biology seeks to quantify changes in biomolecules within tissues and map those changes back to regions, structures and even individual cells within the complex organization of the tissue. A variety of approaches have been developed for measuring mRNA and proteins in tissues and some of these have been developed into commercial platforms. Some commercial platforms determine the gene expression of multicellular regions overlaid on a grid of capture spots arrayed across the tissue section (10x Genomics Visium). Others, such as the Nanostring GeoMx, utilize user-selected regions of interest (ROI) that can specifically target a histologic structure or lesion for expression analysis. For mRNA measurements with these platforms, whole transcriptome scale interrogation of tissues is possible, which is especially suited for discovery applications. Newer platforms such as the 10x Genomics Xenium, Nanostring CosMx, Vizgen MerScope and others have pushed the envelope further and have achieved single-cell resolution in profiling tissues, though not yet at full transcriptome scale.

The rapid pace of deployment of these technologies has sparked interest in applying them to answer clinical questions using patient-sourced tissues. Formalin-fixed paraffin embedded (FFPE) tissue is the workhorse format of anatomic pathology laboratories and is likely to remain so for the foreseeable future. FFPE biospecimens comprise the largest source of patient-derived materials that can be used to understand the pathophysiology of human diseases, develop new diagnostics, and discover new therapeutic interventions. Spatial technologies that can take advantage of this resource have not only the ability to harness the potential of the vast archives of tissue blocks in pathology departments throughout the world, but also have a clear runway for the development of diagnostics tests that can leverage FFPE, the industry standard for processing and stabilizing patient tissue samples for clinical testing (1, 2). Reassuringly, all the leading commercial spatial transcriptomics platforms are compatible with FFPE tissues.

Simultaneous and spatially registered interrogation of multiple biomolecules can provide unprecedented insight into their biological interactions within tissues and cells. While analyzing as many biomolecules as possible in a single assay (high -plex) is desirable, sensitivity of detection and spatial resolution are also important considerations. However, the trade-off between plex (number of genes) and data quality (sensitivity, specificity and resolution) is not well understood for many of the commercial spatial profiling platforms. Increasing plex increases the overall cost of the assay, as does increasing the area of the tissue interrogated. This increased cost may be incurred in terms of the reagents themselves (probes, sequencing required for readout) or experiment execution time. The expense of spatial transcriptomics experiments has been an impediment to benchmarking the rigor and reproducibility of the results that they generate. Reproducibility of results will be an important consideration for clinical translation of these technologies. Successful clinical implementation will also have to balance the cost of these expensive technologies against the information they could provide.

There are now numerous commercially available platforms to perform spatial transcriptomic profiling, and each system has its own advantages and disadvantages. Here, we describe our experience with one multicellular (GeoMx DSP) and one single-cell resolution platform (CosMx SMI), describe their rigor and reproducibility, and our practical experience comparing results within and across platforms. These results highlight important experimental design considerations for using these platforms that affect sensitivity, scope and cost, and which will impact their potential translation to a clinical setting in the future.

## METHODS

### Human nephrectomy tissues (ethics statement and patient consent)

Tissues were collected in deidentified fashion and with informed consent under the University of Washington’s IRB Study Protocol 1297 and in accordance with the Declaration of Helsinki. Fresh human kidney tissues were sourced from patient’s undergoing nephrectomy for removal of kidney tumors. Samples of uninvolved kidney (4 donors), and a portion of the kidney tumor (renal cell carcinoma, RCC) from one of the donors were fixed in 10% neutral buffered formalin for 24-48 hours and then transferred to 70% ethanol for another 24 hours. Samples from all 4 donors (5 pieces of tissue in total) were paraffin embedded into a single multi-tissue block prior to sectioning. These samples were placed such that all 5 tissues could be covered by a 22 × 22 mm coverslip.

### Clinically sourced human kidney biopsy tissues

With approval from the University of Washington’s IRB, electronic health record searches were performed to identify 14 patients with minimal change disease. Kidney biopsies from 3 patients with normal histology that we have reported on previously (3) were retested together with the minimal change disease biopsies using a whole transcriptome based probeset.

### Digital spatial profiling to reduce reagent utilization and to assess technical reproducibility

Four consecutive 5µm thick sections from the kidney multi-tissue block were applied to charged slides and baked at 60°C for 1 hour. After deparaffinization and rehydration, sections were subjected to heat induced antigen retrieval with Tris EDTA, pH 9 for 15 minutes followed by 1µg/ml proteinase K digestion for 15 minutes at 37°C. All 4 slides were hybridized overnight at 37°C with the Cancer Transcriptome Atlas (CTA) probe mix encompassing 1,825 genes. Per Nanostring’s protocol, sections are usually incubated with 250µl of probe mix and coverslipped using a 40×22 mm RNase-free HybriSlip coverslip (ThermoFisher). However, in order to extend the use of the probes, which are the most expensive component of the digital spatial profiling workflow, we utilized 100µl of probe mix per slide and coverslipped sections using a 22×22mm HybriSlip. The following day, all 4 slides were subjected to stringency washes and counterstained with fluorescently labeled antibodies recognizing pan-cytokeratin (Cy3 channel), CD10 (Cy5 channel) as well as DNA (Syto13, FITC channel) (Fig. 1A). One slide was then loaded into the instrument while the others were stored in 2x SSC in the dark at 4°C for staggered collections. On 3 subsequent days, one of the remaining labeled slides was restained with Syto13 for 10 minutes before loading onto the instrument. After loading onto the instrument and scanning, ROI selection was performed, and UV-released barcode probes were collected in staggered fashion into a 96-well collection plate (24 ROI/day; Fig. 1B). After 4 consecutive days of ROI selection, libraries were generated using a SeqCode construction kit from Nanostring. The libraries were sequenced using a NextSeq 2000 P2 flowcell (100 cycle kit). Sequenced barcodes were mapped to genes and ROIs using the Nanostring GeoMx NGS Pipeline, version 2.0.21.

**Figure 1.**
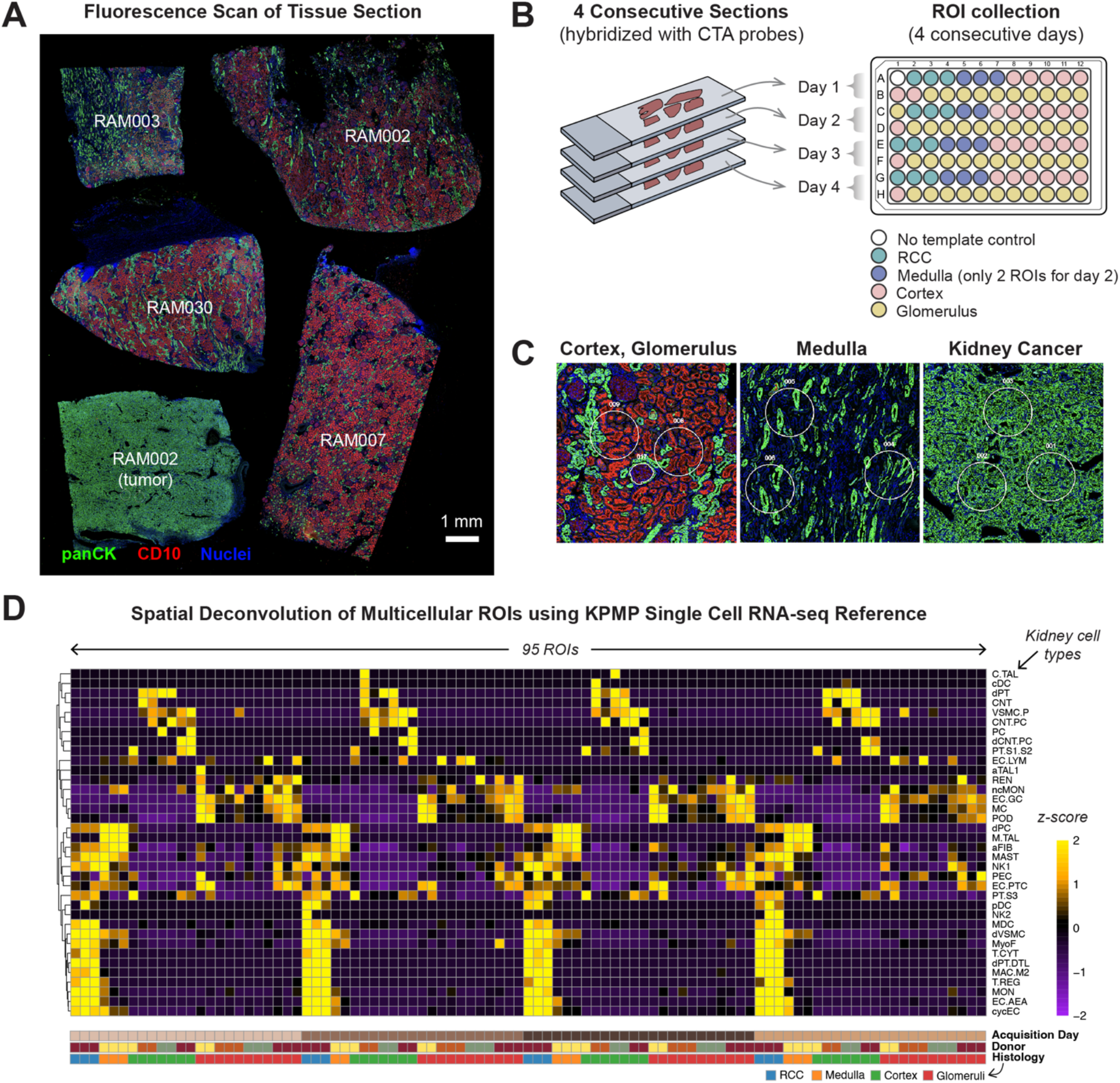
DSP experiment to assess rigor and reproducibility. A) A fluorescence scan from the GeoMx DSP of a section of tissue microarray composed of 4 kidney tissues and 1 kidney cancer tissue from 4 donors. B) Experimental design to assess for technical reproducibility. ROI from different regions were collected into wells of a 96-well plate and gene expression from those ROI was quantified. C) Higher power views of ROIs selected for each of 4 histologically distinct kidney structures. D) Spatial cellular deconvolution of individual ROIs reveals the expected cellular composition as a function of histology.

### Digital spatial profiling on clinically sourced human kidney biopsies

This was performed similar to the procedure described above except for the following modifications. To maximize the number of human kidney biopsies analyzed and extend the use of the Whole Transcriptome Atlas (WTA) probe mix encompassing ∼18,000 genes, 5 µm sections from 3-4 individuals were placed on to each slide used in the experiment. Slides were hybridized with the standard buffer volume of 250µl and covered with 40×22 mm HybriSlips. After glomerular and tubular ROI selection and UV-released probe collection, sequencinglibraries were generated using a SeqCode construction kit. Sequencing-based readout was performed using a NextSeq 2000 P2 flowcell and the data were processed using the GeoMx NGS Pipeline.

### Initial data processing

Analysis of DSP data was performed using the Bioconductor *GeomxTools* package, which was developed by NanoString. After reading the data into *R*, quality control steps were performed to identify problematic probes. For the CTA probe mix, each gene is represented by up to 5 unique probes that recognize different sections of the gene’s mRNA. Probes with a geometric mean <10% of the collapsed gene level estimate (e.g., those probes that don’t contribute much to the overall signal of that gene) were filtered out. In addition, probes with >20% of the observations flagged as outliers were removed. After filtering the probes, probe sets were collapsed to the individual gene level. Multicellular ROIs were deconvolved using the KPMP normal reference single cell RNA-seq data set and the *SpatialDecon* algorithm (4, 5). Since ROIs vary in size, cell count and mRNA content, we assessed the performance of normalization methods such as cyclic loess regression and 3^rd^ quantile (Q3) normalization (recommended by Nanostring).

### Sensitivity comparison: CTA vs. WTA

We compared the performance of the CTA probe mix (∼1,825 genes) against the Whole Transcriptome Atlas probe mix encompassing ∼18,000 genes. We had previously generated CTA measurements from n=12 glomeruli of 3 individuals with histologically normal kidney biopsies. In this study, we performed WTA measurements on n=20 glomeruli from those same 3 individuals. After performing QC and Q3 normalization, we identified a subset of 451 genes that were measured above the limit of quantification (LOQ) in the glomeruli hybridized with CTA probes. We then compared the average counts of these genes (across the 3 donors) against the average Q3 normalized counts generated using the WTA probe mix on the same 3 donors’ kidney biopsy tissues.

### Sensitivity comparison: GeoMx vs. CosMx

Second, we performed single cell transcriptional profiling on the kidney multi-tissue block that we had previously used for reproducibility assessment. We utilized the 1,000 gene universal cell characterization panel on the Nanostring CosMx SMI platform. Tissue morphology was delineated using antibodies reactive for pan-cytokeratin, CD45 and cell membranes (cocktail of anti-β2 microglobulin and anti-CD298 antibodies). We tiled 205 fields of view encompassing all the ROIs selected in our GeoMx experiment that assessed reproducibility. After run completion, the data were automatically pushed to the Nanostring AtoMx cloud-based storage and analysis platform, where cell segmentation, assignment of transcripts to cells and initial calculation of spatial neighborhoods was performed. Semi-supervised cell typing was performed using AtoMx’s InSituType module (6). We used the KPMP normal kidney reference dataset as input and permitted up to 3 unknown cell type assignments in order to account for tumor cell types not represented in KPMP (5). Neighborhoods/spatial niches, top marker genes for cell types and cell-cell proximity analysis were also computed using AtoMx’s in built analysis modules. UMAPs and spatial projections of cell type assignments and neighborhoods were generated from the Seurat object in *R*. In order to select pseudo-ROIs corresponding to the ROIs selected in the GeoMx reproducibility experiment, we utilized the Partek Flow analysis platform. Manual pseudo-ROI selection was guided by spatial neighborhood predictions and attempted to approximate the cell number of the ROIs selected in the GeoMx experiment. If the exact structure was not identifiable or not present in the tissue section, a similar structure in close proximity was chosen. 450 genes were shared between the GeoMx CTA and the 1,000 gene CosMx universal cell characterization panel. Across each pseudo-ROI in the CosMx experiment, transcript counts for these 450 genes were integrated and compared to the integrated collapsing per-gene counts for the corresponding ROI in the GeoMx experiment.

## RESULTS

### Staggered collection of spatial transcriptomic data using the GeoMx DSP platform

A map of the tissue sections used for the spatial transcriptomics experiment is shown in the GeoMx immunofluorescence overview scan (Figure 1A). Replicate sections were successfully hybridized with the reduced amount of CTA probe and the smaller HybriSlip (see Methods for detailed explanation). ROIs encompassing histologically distinct structures (glomeruli, cortex, medulla, tumor) were collected in replicate, from each tissue patch (Figures 1B, C). After sample collection, library construction, sequencing readout and mapping, all ROIs produced transcriptomic data that passed quality control thresholds. After applying GeoMx’s recommended Q3 normalization, the cellular composition of the ROIs was computed using the *SpatialDecon* algorithm. This cellular deconvolution from multicellular GeoMx ROIs revealed the expected composition of cell types within the histologically distinct ROIs. For example, constituent cells of glomeruli, namely podocytes (POD), mesangial cells (MC) and glomerular endothelial cells (EC.GC) were identified only in glomerular ROIs. Tumor cells were not present in the KPMP normal kidney reference dataset. However, cellular deconvolution of the RCC ROIs uncovered diverse immune cell types that were largely restricted to the tumor. The pattern of cellular composition in replicate ROIs was consistent with a given histology and reproducible across the 4 days of staggered sample collection. These results showed that a) reduction of CTA probe volume and hybridization area does not adversely affect data collection; b) cellular deconvolution of histologically distinct ROIs produces the expected cell type composition for a given histology, and that c) staggered collection of replicate samples over 4 days produces qualitatively similar data.

### Quantification of sources of variability

To examine the different sources of variability in the data, we first performed a principal component analysis (PCA), in order to partition variability between samples. In a PCA, similar samples should cluster closely and will be separated from dissimilar samples. For example, in Fig. 2A, left panel, all ROIs of the same histology cluster together, clearly separated from ROIs with different histology. This indicates that the sample histology underlies the largest differences in gene expression among samples with lesser contributions from donor and acquisition day (Fig. 2A middle and right panels). When we restricted PCA plots to a single histology such as Glomeruli or Cortex (FSupplemental figure 1), samples clustered first by patient, and then by ROI, indicating that differences between samples were primarily due to histology, followed by subject, and then ROI. Since consecutive sections were used for data generation, it is conceivable that some of the observed variation in expression is also due to variability in ROI selection. Uniform, circular ROIs were used for cortex, medulla and RCC. By contrast, even though we selected ROIs on the same glomeruli over the 4-day experiment, some glomerular profiles remained relatively invariant (Figure 2B, rows 1 and 2), while others changed noticeably (Figure 2B, rows 3 and 4). Tissues are complex structures that have changing cellular compositions at different levels in serial sections of structures. Thus, understanding the contribution of compositional, intra-histology variability to the measured gene expression variation is an important consideration for experimental design and ROI selection. In order to quantify this further, we plotted the distribution of standard deviations for the log2 transformed, normalized expression of each gene grouped according to histology, donor, ROI and acquisition day (Figure 2C). At the top end of the interquartile range, histology accounted for 84.5% of the gene expression variation, with donor-to-donor variation accounting for 7.3% and ROI selection variability accounting for 6.1%. Day-to-day variability, a non-biological/technical source of expression differences in this experiment, accounted for only 2.0% of variation. These analyses showed that the GeoMx platform exhibited good technical reproducibility in our hands and that the majority of gene expression differences could be attributed to anticipated biological sources of variation such as histology, donor and ROI selection parameters.

**Supplemental Figure 1.**
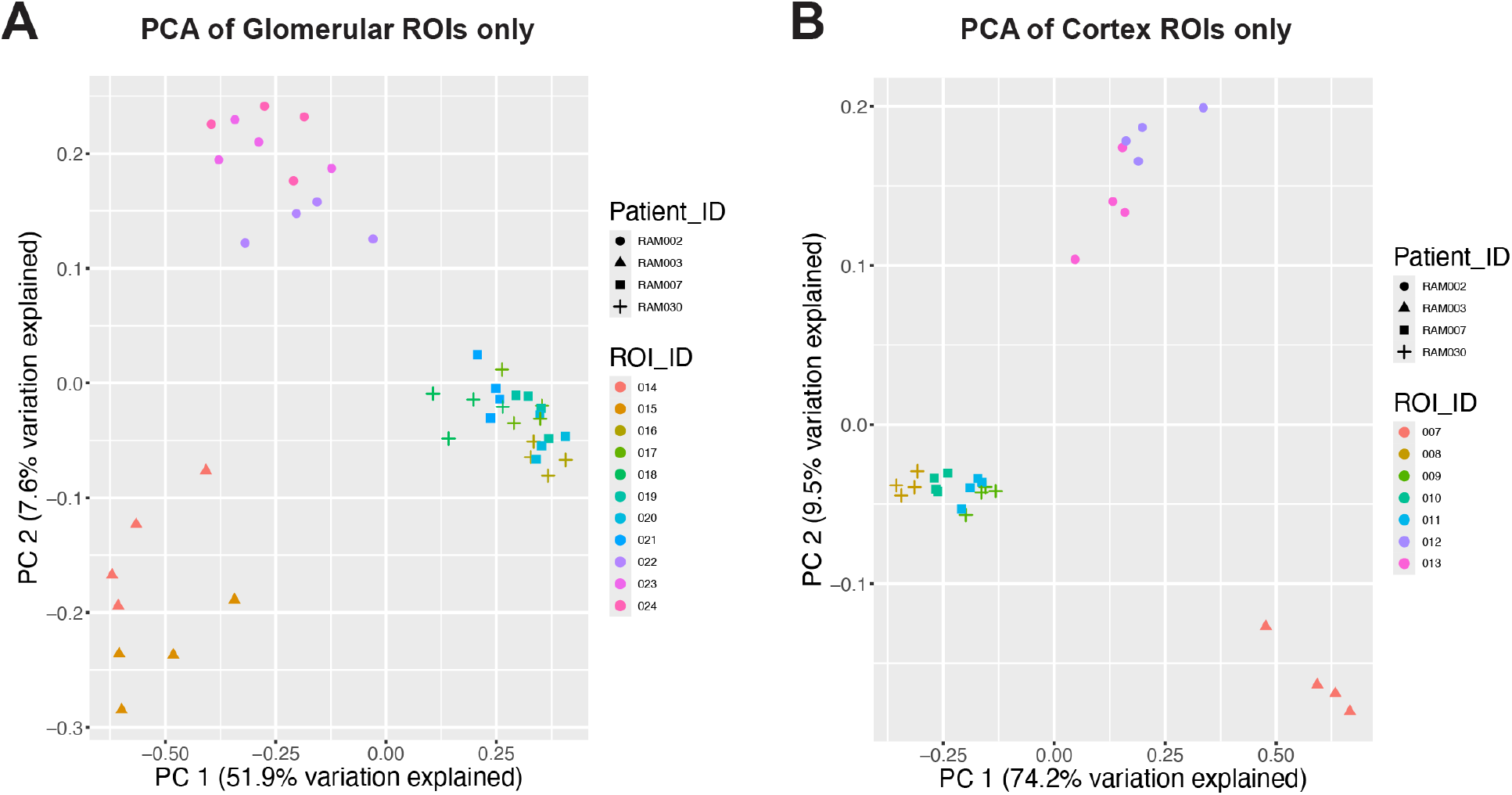
Overview of CosMx 1000-gene experiment performed on kidney multi-tissue block. A) Plot of glomerular ROIs only according to the two greatest principal components (PC1, PC2). A) Plot of cortex ROIs only according to the two principal components (PC1, PC2) contributing the largest variation.

**Figure 2.**
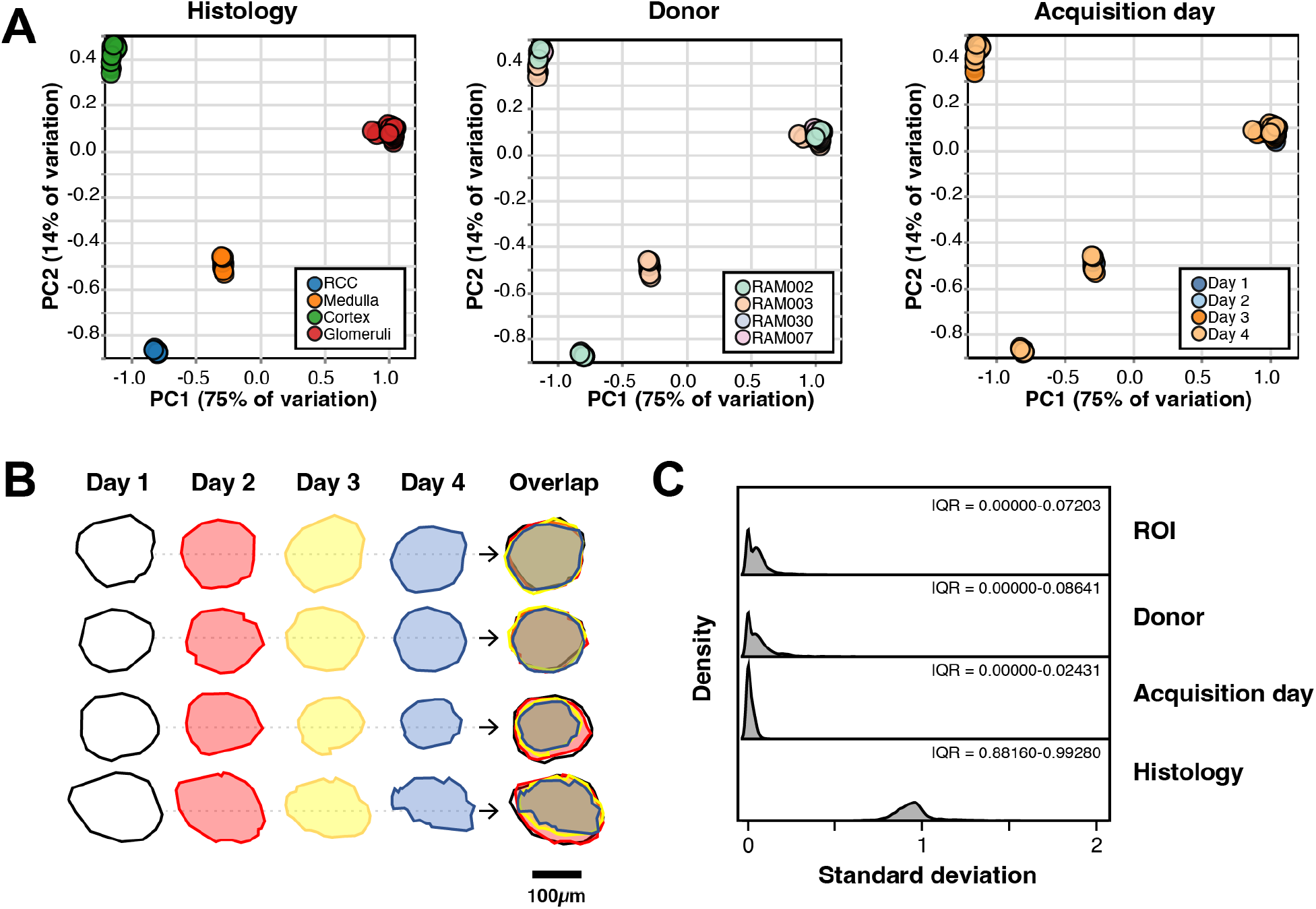
Analysis of sources of variation in DSP workflow. A) Plots of ROIs according to the two greatest principal components (PC1, PC2). Each principal component analysis (PCA) panel is color coded by histology, donor or acquisition day. B) For non-uniform ROIs, the ROI outline introduces another source of variation. When selecting the same glomerulus across 4 consecutive sections, some glomeruli showed relatively invariant profiles (rows 1, 2), while others changed their profiles (rows 3, 4). C) Distribution of standard deviations for measured genes as grouped by histology, acquisition day, donor and ROI.

### Comparison of data normalization approaches

The goal when normalizing data is to remove as much technical variability as possible while retaining biological variability. To do so, we assume that there are one or more genes that have relatively consistent expression across a set of samples, in which case any apparent differences in expression for those genes is primarily technical, and we can therefore use those genes to remove the technical variability via normalization. We used MA-plots to visualize differences between normalization methods. In an MA-plot, the horizontal axis (A) measures the log_2_ average expression level of a gene (so points to the left are lower expressing genes, and those to the right are more highly expressed), and the vertical axis (M) measures the log fold change between the same gene in the two different samples/tissues. Each MA-plot in Figure 3 compares the day 1 sample for a glomerular ROI to a different day. Since these are just different slices from the same tissue, we expect the data to be symmetrically distributed along a horizontal line at zero. This is not the case when no normalization is applied to the data (Figure 3A Unnormalized). The normalization methods that NanoString recommends are simple shift normalizations where either a single observation (Q3 normalization), or the geometric mean of a set of housekeeping genes (negative control normalization) is used to normalize the data. The default normalization recommended by NanoString uses the third quartile (Q3), i.e. 50^th^ to 75^th^ percentile, from each sample to normalize the data. This shift normalization does remove some of the variability but does not account for the fact that the differences between samples varies as a function of gene expression (Figure 3A, Q3 normalized), identified by the red line. Since it is not possible to adjust for these differences using a simple normalization, we next tested methods that account for distributional differences between samples. One such method is called a cyclic loess normalization, which was originally developed for normalization of microarray data (7). The red lines in Figure 3 are locally-weighted regression lines (loess lines) that identify the vertical center of the data at each point on the horizontal axis. As noted above, we assume that there is a set of genes that do not change expression between different samples. We also assume that up and down-regulation is relatively consistent between any two samples. If both assumptions hold, then the loess line identifies the set of unchanging genes, and if we simply adjust the data to linearize the loess line and center it on the horizontal line at zero, we will have removed the technical variability identified by the loess line. We iterate through this process three times (the cyclic part of cyclic loess) to normalize the data. This cyclic loess noticeably improves the log fold change (M) component of the MA plot (Figure 3A, Cyclic loess). We also explored quantile normalization which was reported to be a better method for normalizing GeoMx data between samples with different histology (8). In quantile normalization, observations in each sample are rank-ordered according to magnitude, the average for each rank is computed, and then the average for each rank replaces the observed values. All samples then have identical distributions. Of all the methods tested, quantile normalization produced the most symmetrical distribution along a horizontal line at zero (Figure 3A, Quantile). The different normalization methods make different assumptions of the underlying data structure with expectations for the numbers of differentially expressed genes. These can be visualized using a volcano plot, such as is shown for differential expression between glomerulus and cortex ROIs using the 3 normalization methods (Figure 3B). Examining the overlap of significantly changing genes (FDR<0.05, regardless of magnitude of fold change), the greatest number of differentially expressed genes are identified using Q3 normalization, while cyclic loess and quantile normalization appear to be the more stringent (Figure 3C). Therefore, the choice of normalization methods will affect the downstream analysis and the choice of one method may be dependent on the experimental system and goals. The default Q3 normalization method recommended by Nanostring, appears to be the most permissive and may be more suitable for initial discovery and hypothesis generation.

**Figure 3.**
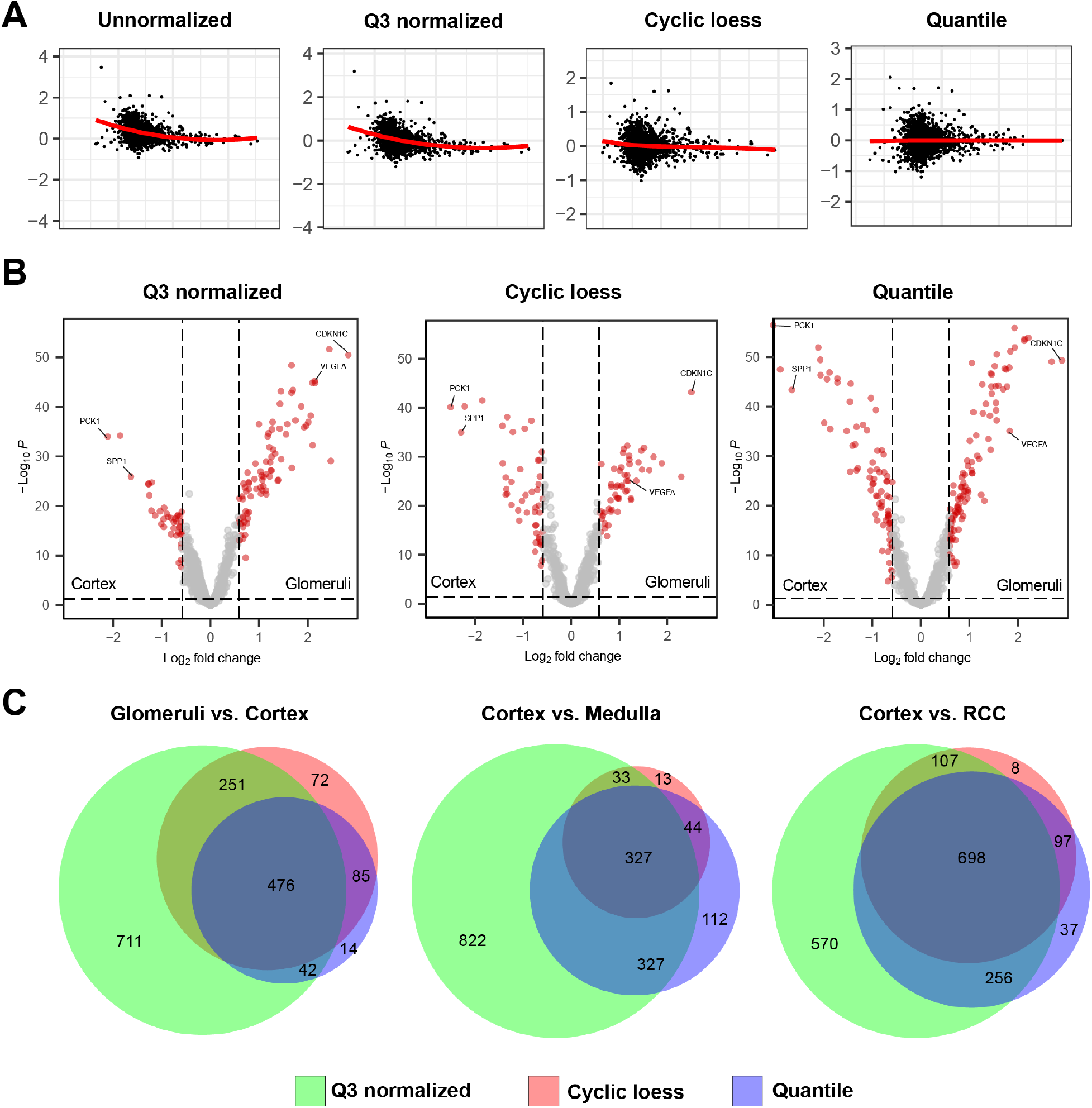
Assessment of normalization approaches for DSP data. A) MA plots for a representative glomerulus showing the impact of various normalization approaches to the distribution of the data. B) Volcano plots showing the impact of normalization approaches on differentially expressed genes detected between cortex and glomerular ROIs (FDR<0.05, fold change >1.5x). C) Venn diagrams of overlap of significantly changing genes (FDR<0.05) in comparison of glomeruli vs. cortex, cortex vs. medulla and cortex vs. RCC.

### Sensitivity comparison: CTA vs. WTA

The reproducibility analysis described above used the CTA probeset, which encompasses 1,825 genes. Each gene within the probeset is detected by up to 5 distinct probes, and the geometric mean of their signals is used to generate the observed expression value. By contrast, in the Whole Transcriptome Atlas (WTA) probeset, each of the >18,000 target genes is detected by only one oligonucleotide reporter. Only 76 of the target genes in the CTA share a probe with the WTA (Kaitlyn LaCourse, Nanostring, *personal communication*), therefore, the majority of the WTA probes are unique to that probe set. This raised two questions: 1) does the CTA have better performance than WTA due to the increased number of probes per target? 2) are expression values generated using WTA comparable to CTA in terms of linearity of response? To assess this, we profiled n=20 glomeruli from n=3 patient biopsy samples using the WTA probe set (Figure 4A). We compared the results to n=12 glomeruli, sampled from those same patients, that were profiled using the CTA in our previous study (3). We identified 451 genes that were above the limit of quantification in our CTA experiment that overlapped with the same targets in the WTA experiment. When we plotted the Q3 normalized counts of these genes using the two probe sets, we observed that there was reasonable linearity of response (r=0.83) (Figure 4B). This indicated that even though different probe sequences were utilized in the CTA and WTA, they still produced comparable results. This analysis also showed that the measured counts using WTA were ∼46% of the CTA, across the target genes in this analysis. This experiment showed that reducing the number of probes/target gene in the hybridization reaction, also reduces the target’s measured expression value.

### Sensitivity comparison: GeoMx vs. CosMx

The GeoMx using a ROI-based strategy to generate spatially localized gene expression profiles. These ROIs are multicellular regions (usually targeting a minimum of 100 cells) and therefore, the resulting expression profiles represent an aggregate of the expression levels of all cells within the ROI. Newer spatial transcriptomics platforms with single-cell resolution (CosMx SMI, Xenium, MERScope) have been released to circumvent this deficiency. In one implementation, the CosMx Spatial Molecular Imager hybridizes gene-specific probes to tissue localized targets followed by rounds of *in situ* detection using fluorescent branched DNA reporters and high-resolution imaging (9). While whole transcriptome detection has been demonstrated, approximately 1,000- or 6,000 genes can be simultaneously profiled using current commercially available probe sets. Importantly, these single-cell platforms use imaging rather than sequencing as their readout modality. They are also more expensive on a per-sample basis than ROI-based profiling approaches. It is therefore necessary to understand the technical tradeoffs of single-cell approaches to understand if their increased cost and resolution is justified in order to answer a research question. To do so, we first performed a 1000-plex CosMx single-cell expression profiling experiment on a section of the multi-tissue block we had previously used for the reproducibility analysis. We tiled 205 fields of view (FOV) to capture gene expression at single-cell resolution across the 5 tissue patches (Supplemental Figure 2A). A total of 284,728 cells were resolved with an average of 132 transcripts detected per cell (10%-90% range = 22 – 274 transcripts/cell). Cloud-based analysis was performed on the AtoMx platform. Semi-supervised cell-type assignment and clustering was performed using AtoMx’s InSituType module (6) (Supplemental Figure 2B). We used the KPMP normal kidney reference dataset as input and permitted up to 3 unknown cell type assignments in order to account for tumor cell types not represented in KPMP (5). Two of the unknown cell types (b, c) were almost exclusively localized to distinct neighborhood comprising the tumor specimen (Supplemental Figure 2C, D), while the third unknown cell type (a) represented a form of tubule cell/state that was not present in the KPMP normal cell atlas.

**Supplemental Figure 2.**
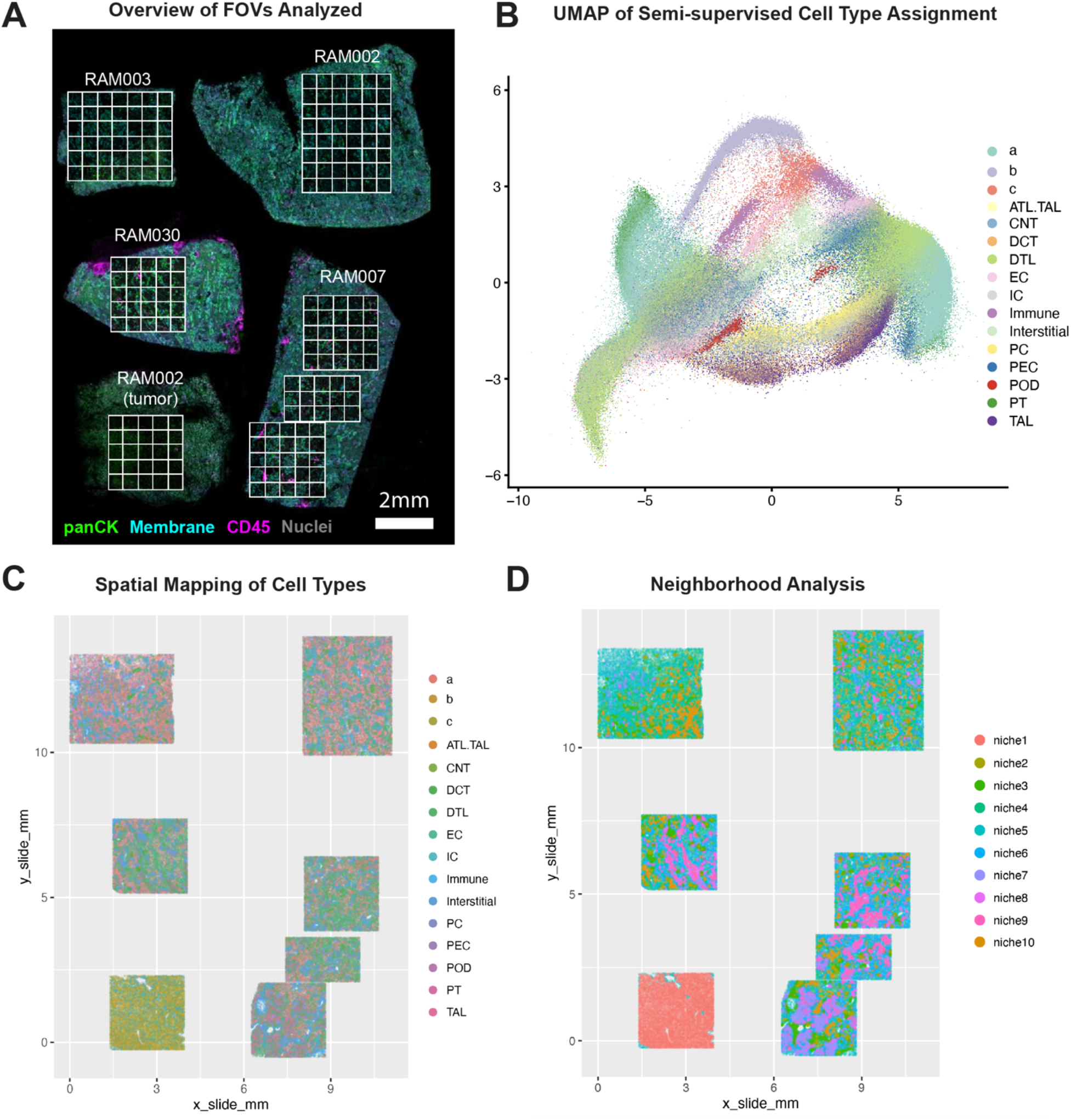
Overview of CosMx 1000-gene experiment performed on kidney multi-tissue block. A) Fluorescence preview scan of tissue section with boxes indicating fields of view (FOVs) from which 1000-gene expression information was captured at single-cell resolution. B) UMAP of imputed cell types from the experiment, including known cell types from the KPMP reference dataset as well as 3 unknown cell types. C) Spatial mapping of imputed cell types back to tissue locations. D) Neighborhood/niche analysis of cell type organization reveals distinct structures in normal kidney tissues and a relatively homogenous neighborhood in this portion of RCC.

A heatmap of expression of the top marker genes for each cell type identified the expected pattern of expression for known kidney cell types. Unknown cell type b likely corresponded to RCC tumor epithelial cells since they expressed *CLU, LDHA* and *CD24* (10-12) (Supplemental Figure 3A). Unknown cell type c likely represented tumor endothelial cells, since they expressed *IGFBP3* and *VW*, which we have recently characterized as RCC tumor endothelial cell selective marker genes in an orthogonal single cell RNA-seq study using different patient tissue samples (13).

**Supplemental Figure 3.**
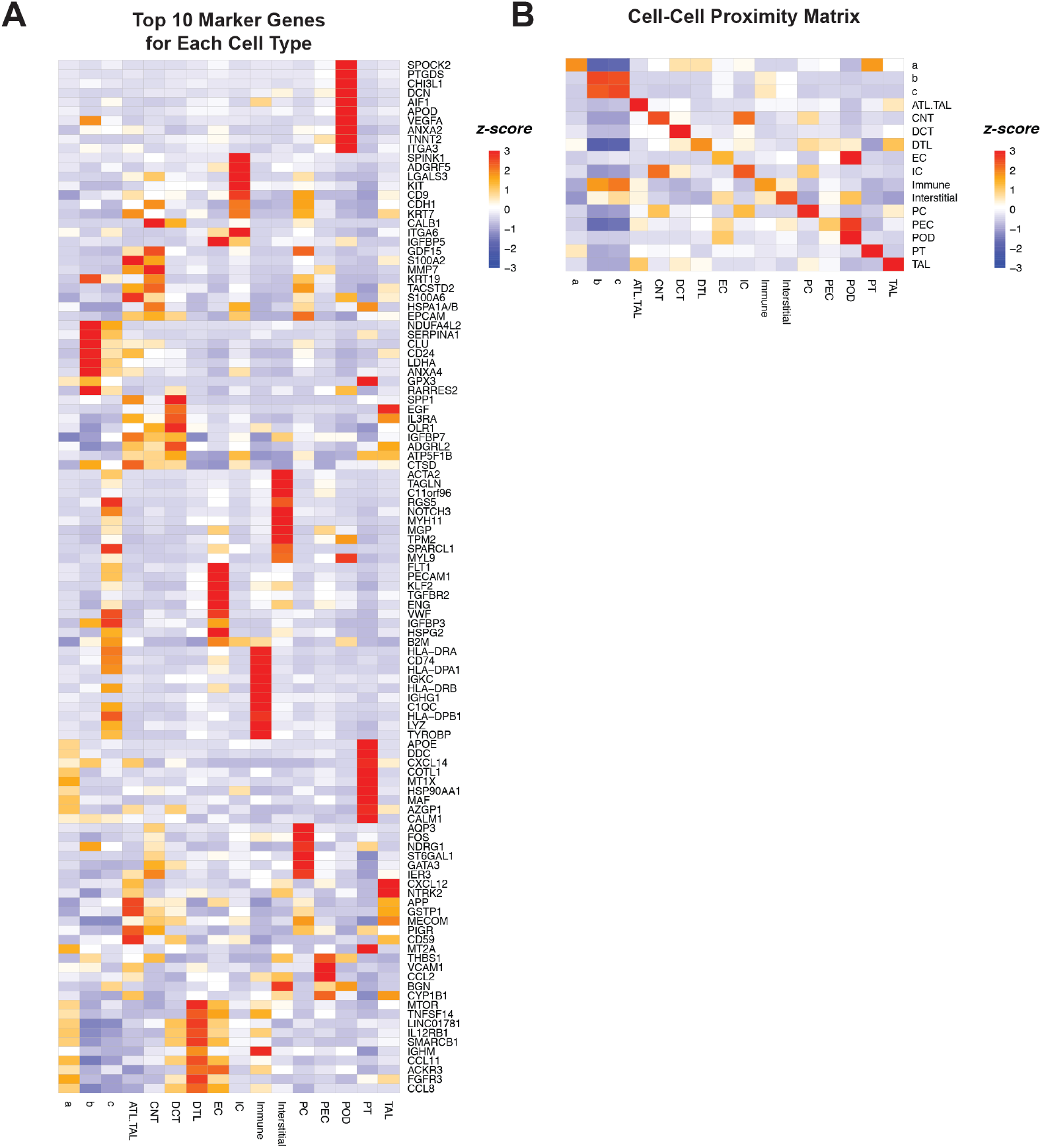
Prediction of marker genes and cell-cell proximity analysis. A) Heatmap of top 10 marker genes for each cell type. B) Proximity analysis of cell types from CosMx single-cell data set performed on human kidney tissues.

These initial analyses gave us confidence that the CosMx platform was capable of recapitulating key geographic and transcriptional features of kidney tissue. To determine if the sensitivity of detection was different for the CosMx platform compared to GeoMx, we defined pseudo-ROIs in that dataset to match the ROIs selected in the GeoMx experiment (Figure 4C). The number of cells in the GeoMx ROIs was comparable to those in the CosMx pseudo-ROIs (Figure 4D). We integrated counts for the 450 genes present in both datasets (GeoMx Cancer Transcriptome Atlas, CosMx 1K Discovery Panel) and found that there was good correlation (r=0.88) of expression levels across both datasets (Figure 4E). Similar to the CTA, the CosMx 1K Discovery Panel uses 5 probes to detect each gene (Kaitlyn LaCourse, Nanostring, *personal communication*). Even so, the integrated counts for the CosMx pseudo-ROIs were 58% of the GeoMx ROIs. Therefore, even though the GeoMx platform uses a sequencing-based readout and the CosMx platform uses an imaging-based readout, both produce comparable patterns of gene expression, though there is reduced signal when single-cell resolution is desired.

**Figure 4.**
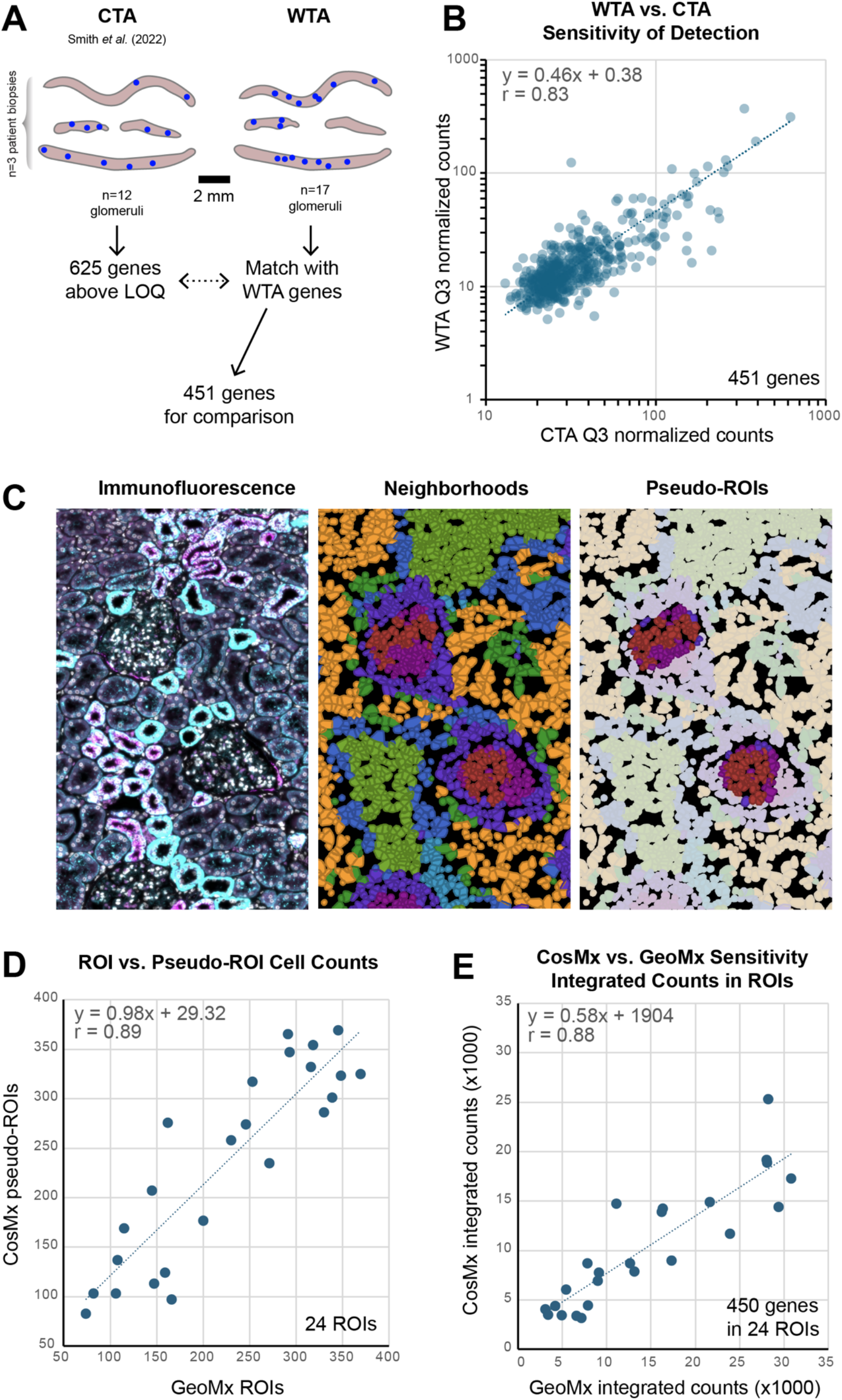
Comparison of CTA vs. WTA and GeoMx vs. CosMx. A) Overview of comparison of glomerular gene expression from n=3 normal kidney biopsies (middle biopsy had two fragments). Blue dots indicate the location of the glomeruli profiled in each experiment. B) Dot plot of Q3 normalized counts of genes measured in glomeruli from kidney biopsies using CTA vs. WTA. C) *Left*, Immunofluorescence overview of a portion of kidney tissue profiles using the CosMx platform and a 1000-gene panel. *Middle*, the expression data was processed to infer cell type and cellular neighborhoods. *Right*, individual cells within histologic structures were grouped together as pseudo-ROI in order to correspond to the ROI chosen in the reproducibility experiment (compare to Figure 1C). D) Plot of cell counts in CosMx pseudo-ROIs vs. cell counts based on nuclear staining in corresponding GeoMx ROIs. E) Correlation of integrated gene counts in CosMx pseudo-ROIs vs. their corresponding GeoMx ROIs.

## DISCUSSION

There is increasing emphasis on improving the replicability of scientific studies, in grant applications, publications and clinical trials (14-16). Understanding how experimental procedures and technologies influence the ability to produce rigorous and reliable data forms the bedrock of replicability. Determining the technical requirements for generating rigorous and reproducible spatial transcriptomics data is critical to align these expensive technologies with experimental questions. It will also form the basis for making economic decisions when planning research studies and for determining which spatial transcriptomics technologies may be amenable to clinical translation. In this study we demonstrate that altering the hybridization procedure for the GeoMx platform allows the researcher to extend the probe set to more slides, increasing flexibility in experimental design, and decreasing the per slide cost. Our results also demonstrate that the greatest source of variability is histology, which is the desired experimental variable. Inter donor variation is the next greatest source of variation, followed by variation of feature (ROI) selection across multiple sections. Our results also demonstrate that it is better to prioritize collection of biological replicates over technical replicates. Reassuringly, day-to-day variation was the smallest contributor to the observed differences. These findings demonstrate that the GeoMx platform allows for flexible spatial interrogation of transcriptomes with relatively little technical noise, allowing for the identification of biological differences in gene expression such as those between histologically distinct structures and among donors.

When analyzing transcriptomic data from the GeoMx, the recommended normalization procedure is Q3 normalization. We found that Q3 normalization of WTA data was relatively permissive compared to alternative methods such as cyclic loess and quantile normalization. The Venn diagrams in Figure 3C indicate that Q3 normalization is likely to identify more candidate differentially expressed genes (DEGs) than cyclic loess or quantile normalization, but they also show that that DEGs identified with the different normalization methods may be distinct. It is important to realize that different normalization methods will influence results and identification of DEGs, and selection of the normalization method may be influenced by experimental goals. The platform’s recommended Q3 normalization may be sufficient when trying to identify the greatest number of DEGs in pilot and feasibility studies, and more rigorous normalization approaches may be more suited for identification of candidate genes for development of a multiparameter disease classifier.

Within the GeoMx platform, there are 2 off-the-shelf probe sets for use with human tissues, the CTA and the WTA. Since GeoMx probes are designed with a UV-cleavable linker and reporter oligonucleotide, expanding the target space from CTA (1,825 genes) to WTA (>18,000 genes) incurs a significant probe synthesis cost. This is somewhat mitigated by reducing the number of the probes/target gene from 5 probes/gene in CTA to 1 probe/gene in the WTA probe set. However, when we compared DEG for the CTA and WTA probe sets on the same set of kidney biopsies, we found that while there was good correlation in expression trends, the WTA was half as sensitive as the CTA. Thus, probe set choice can also influence experimental results, and increasing plex will come at the cost of decreasing sensitivity and the ability to detect low expressing genes. This also suggests that implementing a GeoMx workflow in the clinic would benefit from a smaller number of target genes being analyzed, but with more probes/target to maximize sensitivity. Initial exploratory analyses using the WTA could be used to define the gene subset that is maximally informative for the clinical question at hand before synthesizing the targeted panel of probes.

The CosMx platform permits subcellular resolution of spatial transcriptomic data. Although computational cellular deconvolution strategies can estimate cell type abundance within regions of interest, the higher resolution of CosMx allows for more precise identification of cell types, as well as the identification of novel cell types and cell states. The relationships between cells, cellular neighborhoods, can also be defined with CosMx data. Our analysis confirmed that CosMx could properly identify many of the known kidney cell types that have been identified in the KPMP, as well as to predict novel cell types that are only found in tumor tissue. Gene expression between the GeoMx CTA and the CosMx 1000-plex gene panel showed good correlation (R=0.88) for the 450 genes that were detected on both platforms in spite of their different methods of readout (sequencing vs. imaging). Therefore, the somewhat reduced sensitivity and increased cost/sample of single-cell resolution platforms such as CosMx must be balanced against its increased resolution to provide biological insight into tissue physiology and disease processes. These factors will also dictate which clinical scenarios will benefit from CosMx’s single-cell resolution vs. the multi-cellular resolution afforded by GeoMx.

In summary, we demonstrate that the GeoMx and CosMx spatial transcriptomics platforms are robust tools for spatially registered gene expression analysis. The GeoMx platform lends itself to greater flexibility and potential cost savings by varying the hybridization area and probe set design and volume. Choices for data normalization for GeoMx will affect results and the detection of DEGs. The potential sources of noise (donor-to-donor variation, day-to-day variation in ROI selection, and intrasample ROI selection of biologically similar structures) are small compared to the biological variation that can be measured when selecting histologically distinct structures (e.g. glomeruli, cortex, medulla and RCC). Probe set composition (number of probes/target gene) as well as platform methodology (sequencing readout vs. imaging) can also impact sensitivity of detection. Understanding how these technical and platform considerations influence experimental results will help guide researchers through the planning and execution stage of spatial transcriptomic experiments and facilitate their adoption to answer clinical questions.

## AUTHOR CONTRIBUTIONS

SA and KDS conceived of the study. XL, GS and EB performed spatial transcriptomics experiments. SA, KDS, JWM and TKB performed data analysis and visualizations. SA and KDS composed the original manuscript draft and figures. All authors read, edited and approved the final version of the paper.

